# Tissue-specific endothelial cell heterogeneity in human type 2 diabetes

**DOI:** 10.64898/2025.12.01.691646

**Authors:** Alexander-Francisco Bruns, Oliver I Brown, Hema Viswambharan, Klaus K Witte, Sam Straw, John Gierula, Chew Weng Cheng, Katherine Bridge, Natalie Haywood, Anna Skromna, Natallia Makava, Lauren F Daly, Matthew Bourn, Jeanne F Rivera, Nora Almuhim, Michael Drozd, David J Beech, Lee D Roberts, Khalid Naseem, Richard M Cubbon, Mark T Kearney

**Author notes:** Address for Correspondence: Professor Mark Kearney, The LIGHT Laboratories, University of Leeds, Clarendon Way, Leeds, LS29JT, United Kingdom,. Denotes equal contribution (Alexander-Francisco Bruns, Oliver I Brown).

## Abstract

Transcriptomic heterogeneity of microvascular endothelial cells (MVEC) within subcutaneous adipose tissue (SAT) in preclinical models of obesity/type 2 diabetes mellitus (T2DM) is well-established. Here, we examined if such heterogeneity exists between two canonical insulin-target tissues, SAT and skeletal muscle (SkM). Characterisation of SAT and SkM MVEC from humans with T2DM demonstrated that SAT, but not SkM, exhibited cellular stress at transcriptomic, morphological, and protein levels.

## Main

Dysfunction of the microvasculature is essential in the pathogenesis of T2DM induced tissue damage (1,2) due to its role in regulating perfusion and metabolic exchange between the circulation and tissues (3). The distinctive vulnerability of SAT and SkM in T2DM is well established (4). To shed light on the yet unclear mechanisms underpinning this phenomenon, we explored the possibility that MVEC demonstrate heterogeneous responses to T2DM in SAT and SkM.

We collected pectoral SAT and SkM samples from 61 individuals with and without T2DM undergoing implantation of a cardiac electronic implantable devices (**see Supplementary Table 1 for details**). MVEC were isolated (5) from SAT and SkM achieving ∼99% purity (**Supplementary Figure 1a-c**). We found no evidence of endothelial to mesenchymal transition (**Supplementary Figure 1d—f**) or senescence (**Supplementary Figure 2**). Unbiased bulk RNA sequencing revealed 52 differentially expressed genes (DEG) in T2DM compared with ND SAT MVEC (**Figure 1a,b**) and 3 in SkM MVEC (all P<0.05) (**Figure 1c,d**). Gene ontology (GO) enrichment analysis using g:Profiler highlighted terms representing proteolysis (GO Term 0030162), cellular stress (GO Term 0033554) and endothelial cell migration (GO Term 0010596) in SAT MVEC (**Figure 1e**). Perturbations in cellular morphology are thought to reflect increased cell stress (6,7), and image-based profiling (8,9) depicted by Leiden clustering analysis revealed the presence of morphological differences in SAT MVEC not seen in SkM MVEC (**Figure 1f—h**). Based on these results, we next examined presence of endoplasmic reticulum stress (ERS), a pivotal process in the pathophysiology of T2DM (10,11). T2DM SAT MVEC showed significantly increased levels of the transcription factor C/EBP homologous protein (CHOP), its downstream targets interleukin-6 (IL-6) and interleukin-8 (IL-8), as well as decreased levels of endothelial nitric oxide synthase (eNOS) (12—14) (**Figure 2a,b**). We observed no difference in other inflammatory mediators (**Supplementary Figure 3**), other proteins relevant in ERS (**Supplementary Figure 4**), or markers of autophagy (**Supplementary Figure 5**) in MVEC from either tissue. Correlation and regression analysis of CHOP with clinical variables demonstrated a positive relationship between CHOP and Haemoglobin A1c suggesting a clinically relevant link between CHOP expression and glucose control in SAT MVEC (**Figure 2c,e**) but not SkM MVEC (**Figure 2d**).

**Figure 1.**
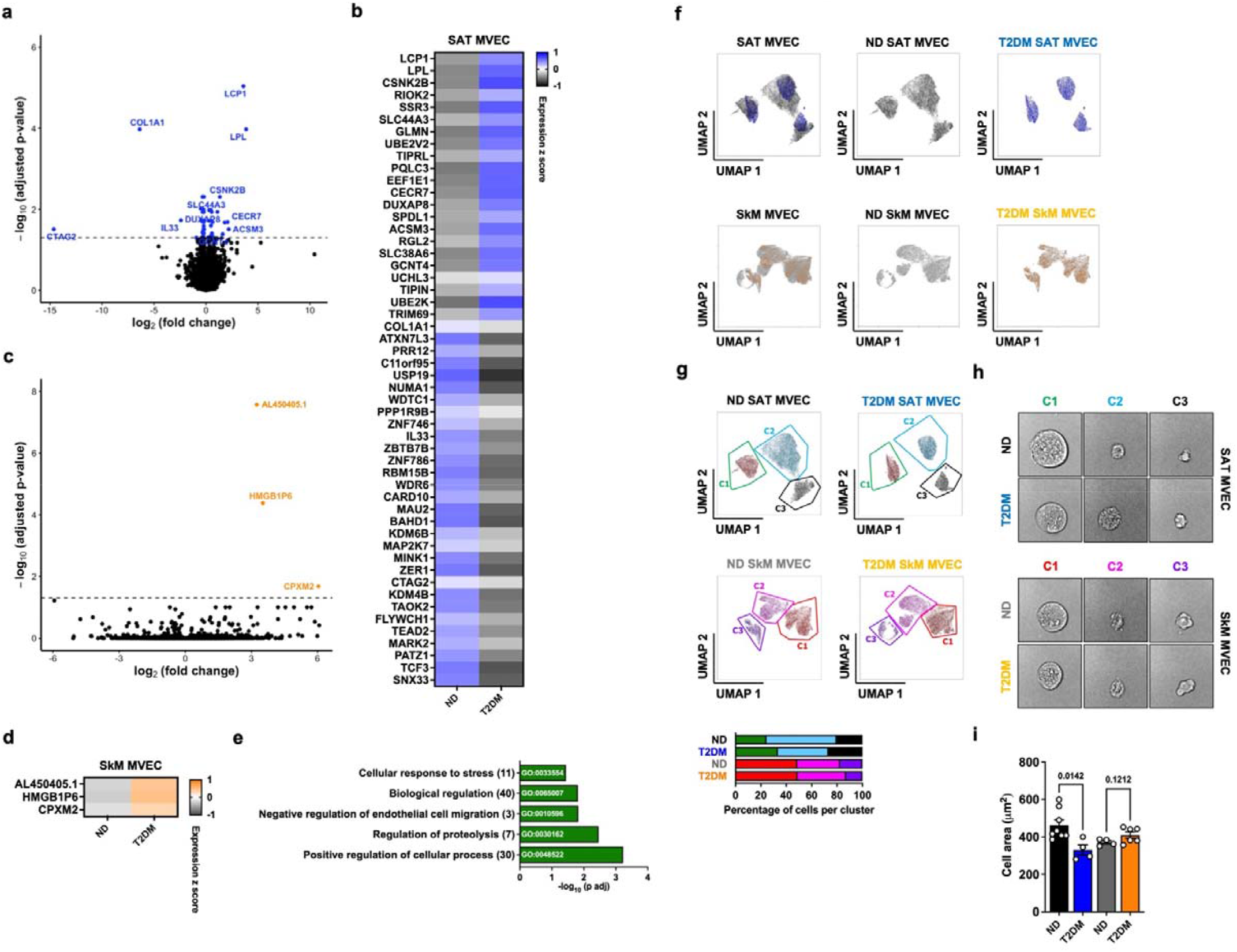
Heterogeneity of SAT and SkM MVEC. **a, c**, Volcano plots comparing differentially expressed genes (DEG) from (**a**) SAT MVEC isolated from non-diabetic (ND, n=25) and T2DM (n=17) individuals or (**c**) SkM MVEC from ND (n=16) and T2DM (n=9) individuals. Significant DEG (adjusted p-value ≤0.05) are shown in blue (T2DM SAT MVEC) or orange (T2DM SkM MVEC). DEG with log2 fold change of –1 or +1 are highlighted. **b, d**, Expression z-score of DEGs in SAT MVEC (**b**) and SkM MVEC (**d**). Grey represents reduced expression, blue or orange increased expression. **e**, Five selected gene ontology (GO) terms from SAT MVEC are shown. In brackets gene count, x-axis represents −log10 of adjusted p-value. **f**, UMAP clustering of SAT MVEC (ND, n=8; T2DM, n=4) and SkM MVEC (ND, n=4; T2DM, n=6) based on cell size showing overlap of clusters. **g**, Highlighting of distinct clusters shown in (**f**). Quantification of clusters shown; the same colour code has been used. **h**, Example images of cells from each cluster shown in (**g**). **i**, Quantification of cell area of SAT and SkM MVEC used in (**g**). Data are presented as mean □ ±□ SEM and were tested for normal distribution prior to analysis by two-tailed unpaired Student’s *t*-test or Mann-Whitney test as appropriate.

**Figure 2.**
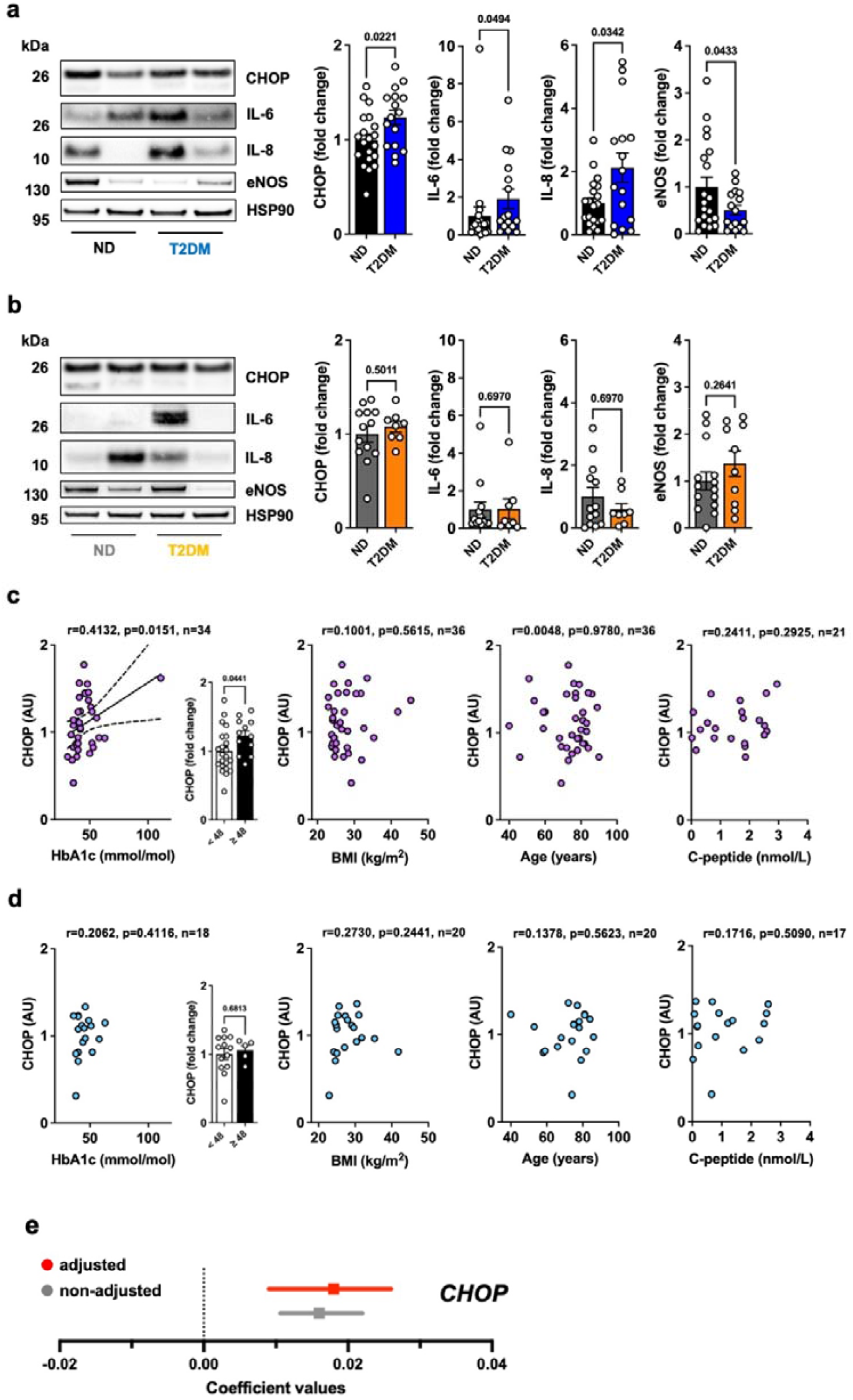
Presence of ER-stress is correlated with HbA1c-levels in SAT MVEC. **a**, Representative Western blots and quantifications showing upregulation of protein expression of CHOP, IL-6, IL-8, and downregulation of eNOS (n=20 ND/ 16 T2DM) in T2DM SAT MVEC, but not in (**b)** T2DM SkM MVEC: CHOP, IL-6, IL-8 (n=13/ 8), eNOS (n=14/ 10). Lysates were run on separate gels each with their own loading control, blots were processed under identical conditions and cropped for clarity. **c, d**, Correlations for CHOP and HbA1c, BMI, age, or C-peptide in SAT MVEC (**c**) and SkM MVEC (**d**). Where significant, linear regression with 95% confidence intervals is shown, n= number of pairs. Bar charts represent CHOP expression in each group with a HbA1c-level of ^3^48 considered T2DM. In bar charts, data are presented as mean □ ± □ SEM. All data were tested for normal distribution prior to analysis by two-tailed unpaired Student’s *t*-test, Mann-Whitney test (**a** and **b**), Spearman or Pearson correlation (**c** and **d**) as appropriate. **e**, HbA1c regression coefficient for CHOP adjusted for age, sex, and BMI. Data is presented with 95% CI.

A recent comprehensive study showed obesity-driven disruption of EC gene expression in an organ-specific fashion in mice (15). A major advance in creating a single-cell transcriptome atlas of human SAT EC has recently been made (16). However, to our knowledge we are the first to demonstrate a link between glycaemic control and CHOP expression in human SAT MVEC and tissue (subcutaneous adipose vs skeletal muscle) selectivity of this potentially deleterious response.

In conclusion, our data suggest heterogeneity of MVEC in 2 canonical insulin target tissues from humans with advanced T2DM. Furthermore, SAT MVEC from humans with T2DM are particularly vulnerable to the deleterious environment created by T2DM resulting in a selective increase in expression of the proinflammatory/proapoptotic transcription factor CHOP. Accordingly, we found alterations of IL-6, IL-8, and eNOS at a protein level. We further demonstrated a significant relationship between poor glycaemic control (as evidenced by elevated glycosylated haemoglobin) and CHOP. This suggests that deliberate and early targeting of glycaemic control or ER-stress could alleviate the cardiovascular risk associated with T2DM.

## Acknowledgements

This research was supported financially by the British Heart Foundation (BHF) Accelerator Award (RE/23/130040); BHF Program Grant RG/F/22/110076 and National Institute for Health and Care Research (NIHR including the NIHR Leeds Biomedical Research Centre NIHR203331. The views expressed are those of the authors and not necessarily those of the NHS, the NIHR or the Department of Health and Social Care.

## Author contributions

M.T.K., K.K.W. and D.J.B. conceptualized the study. A.F.B., OIB., H.V., A.S., N.M., K.P., N.H., M.B., L.F.D., J.F.R., and N.A., designed and performed the experiments. R.M.C, A.F.B., O.I.B., C.W.C., H.V., and M.D. analysed and interpreted data. M.T.K. and A.F.B. drafted the manuscript. N.N., K.N, O.I.B., K.P., S.S., D.J.B., L.D.R. and R.M.C. edited the manuscript and provided important intellectual content. All authors approved the final version of the paper.

## Competing interests

The other authors declare no competing interests. All authors declare that this work was not influenced by economic or other types of conflicting interest.

---

**Supplementary Figure 1.**
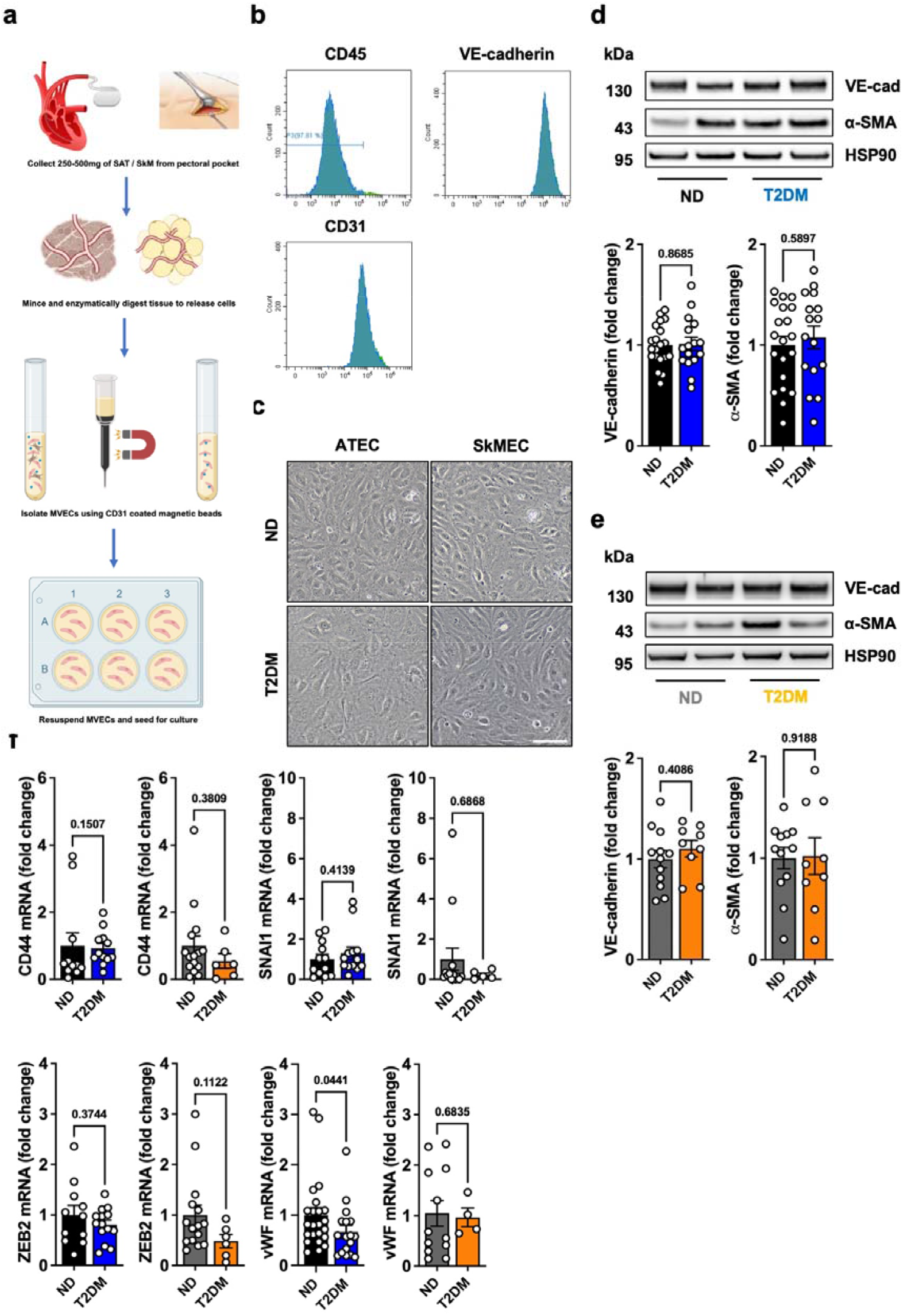
T2DM SAT and SkM MVEC do not display EndMT. **a**, Schematic workflow of MVEC isolation from SAT and SkM. **b**, Representative flow cytometry results from cultured MVEC. Cells routinely showed ∼99% purity as determined by presence of EC markers VE-cadherin and CD31, and absence of CD45. **c**, Representative images of cultured SAT MVEC and SkM MVEC at confluency. Scale bar = 100 μM. **d, e**, Representative blots and quantifications showing expression of VE-cadherin and α-SMA in n=20 ND/ 16 T2DM SAT MVEC (**d**) and n=12 ND/ 9 T2DM SkM MVEC (**e**). **f**, quantifications of RT-qPCR data for genes involved in EndMT: CD44 (SAT MVEC, n=11 ND/ 12 T2DM; SkM MVEC, n=15 ND/ 6 T2DM), SNAI1 (n=12 ND/ 13 T2DM; n=15 ND/ 5 T2DM), ZEB2 (n=11 ND/ 14 T2DM; n=15 ND/ 6 T2DM), vWF (n=23 ND/ 17 T2DM; n=12 ND/ 4 T2DM). In bar charts, data are presented as mean ± SEM. All data were tested for normal distribution prior to analysis by two-tailed unpaired Student’s *t*-test or Mann-Whitney test (**d, e**, and **f**) as appropriate.

**Supplementary Figure 2.**
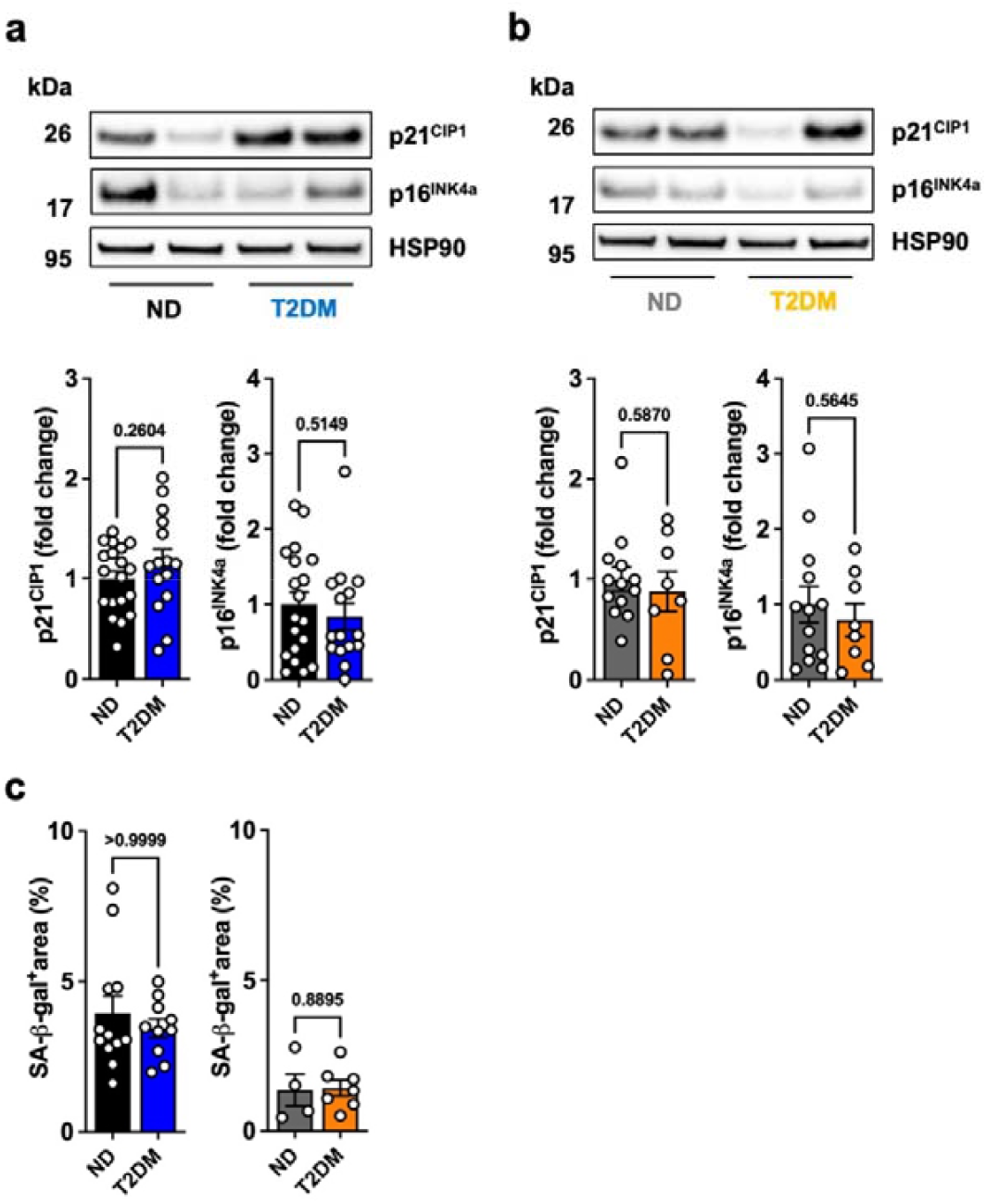
T2DM SAT and SkM MVEC do not display senescence. **a, b**, Representative blots and quantifications of cell cycle proteins p21 and p16 from cultured (**a**) SAT MVEC (n=19 ND/ 15 T2DM), or (**b**) SkM MVEC (n=13 ND/ 8 T2DM). **c**, Quantification of % SA-β-gal positive area in cultured SAT MVEC (n=12 ND/ 10 T2DM) and SkM MVEC (n=4 ND/ 7 T2DM) at confluency. All data are presented as mean ± SEM and were tested for normal distribution prior to analysis by two-tailed unpaired Student’s *t*-test or Mann-Whitney test as appropriate.

**Supplementary Figure 3.**
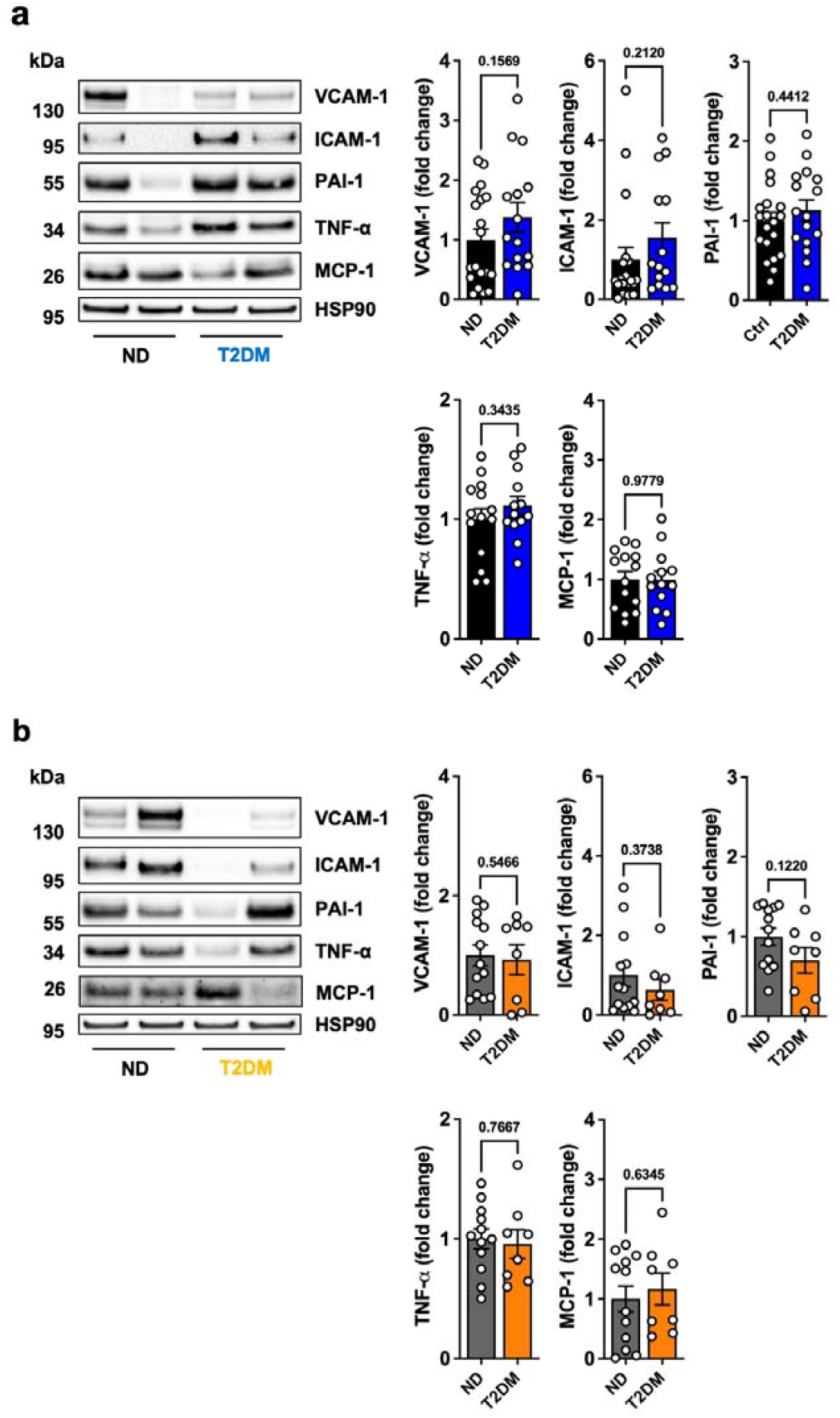
T2DM SAT and SkM MVEC have no inflammatory proteins. **a, b**, Representative blots and quantifications of inflammatory proteins (**a**) SAT MVEC: VCAM-1 (n= 19 ND/ 15 T2DM), ICAM-1 (n=19 ND/ 14 T2DM), PAI-1 (n=20 ND/ 16 T2DM), TNF-α, and MCP-1 (n=14 ND/ 13 T2DM, each). (**b**) From SkM MVEC: VCAM-1, ICAM-1, PAI-1 (n=13 ND/ 8 T2DM), TNF-α, and MCP-1 (n=12 ND/ 8 T2DM). Data are presented as mean ± SEM. All data were tested for normal distribution prior to analysis by two-tailed unpaired Student’s *t*-test or Mann-Whitney test as appropriate.

**Supplementary Figure 4.**
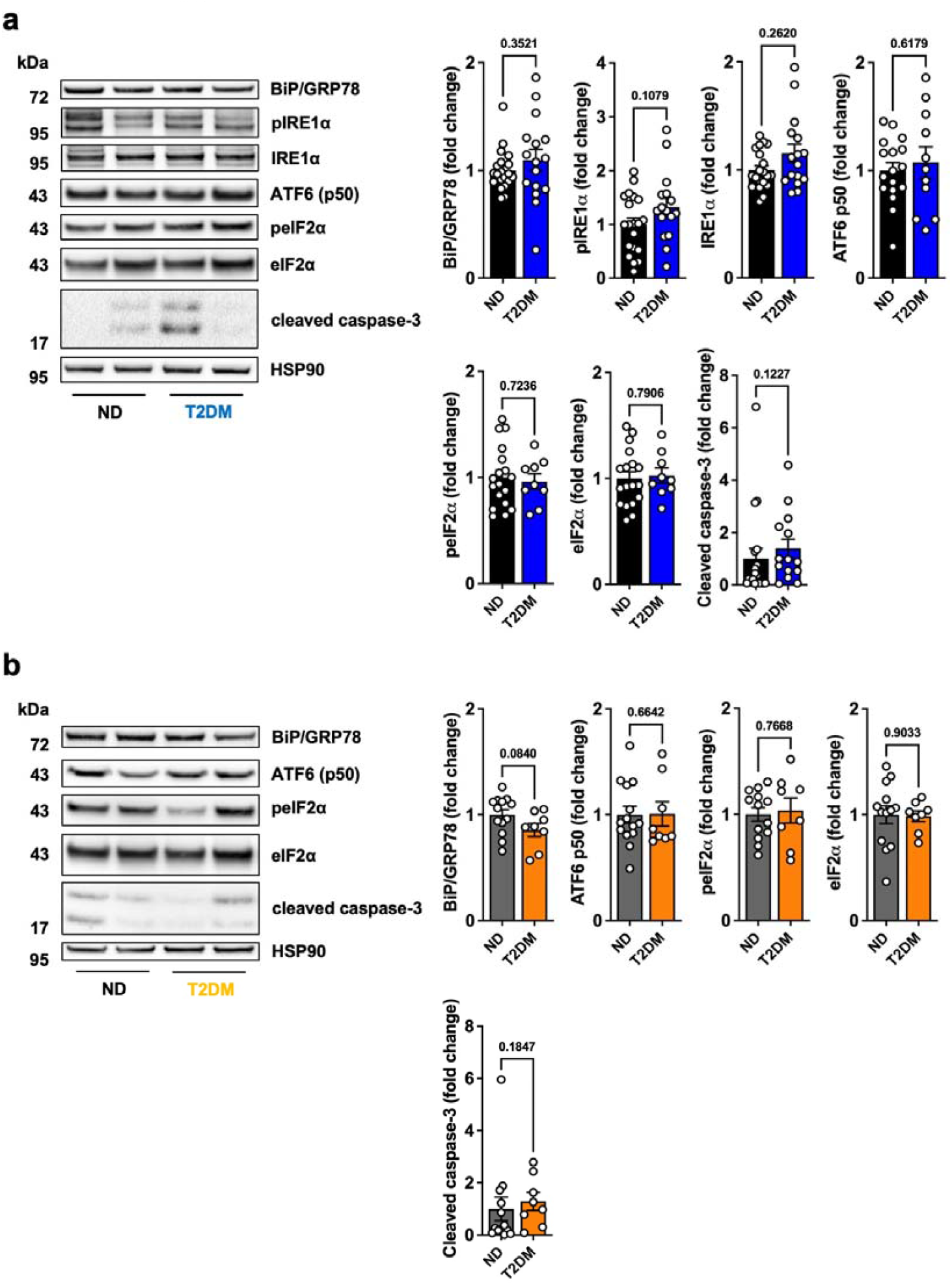
Markers of ER-stress and SkM MVEC. **a, b**, Representative blots and quantifications of proteins involved in the unfolded protein response from (**a**) SAT MVEC: BiP/ GRP78, phospho and total IRE1α (n= 20 ND/ 16 T2DM), ATF6 (n=17 ND/ 11 T2DM), phospho and total eIF2α (n=18 ND/ 9 T2DM), cleaved caspase-3 (n=19 ND/ 14 T2DM). (**b**) From SkM MVEC: BiP/ GRP78 (n= 13 ND/ 8 T2DM), ATF6 (n=14 ND/ 8 T2DM), phospho and total eIF2α as well as cleaved caspase-3 (n=13 ND/ 8 T2DM). Data are presented as mean ± SEM. All data were tested for normal distribution prior to analysis by two-tailed unpaired Student’s *t*-test or Mann-Whitney test as appropriate.

**Supplementary Figure 5.**
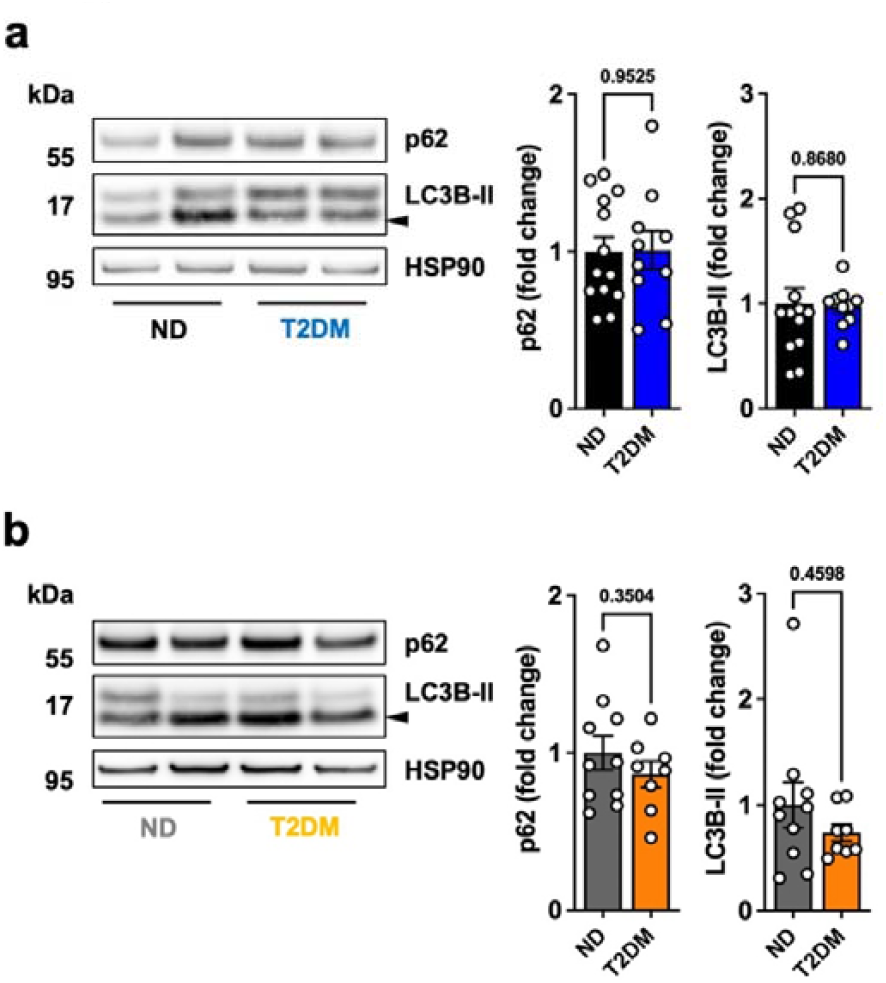
T2DM SAT AND SkM MVEC maintain noraml level of autophagy. **a, b**, Representative blots and quantifications of proteins involved in autophagy from (**a**) SAT MVEC: p62 and LC3B (n=13 ND/ 10 T2DM), (**b**) from SkM MVEC: p62 and LC3B (n=10 ND/ 8 T2DM). Data are presented as mean ± SEM. All data were tested for normal distribution prior to analysis by two-tailed unpaired Student’s *t*-test or Mann-Whitney test as appropriate.

**Supplementary Table 1.**
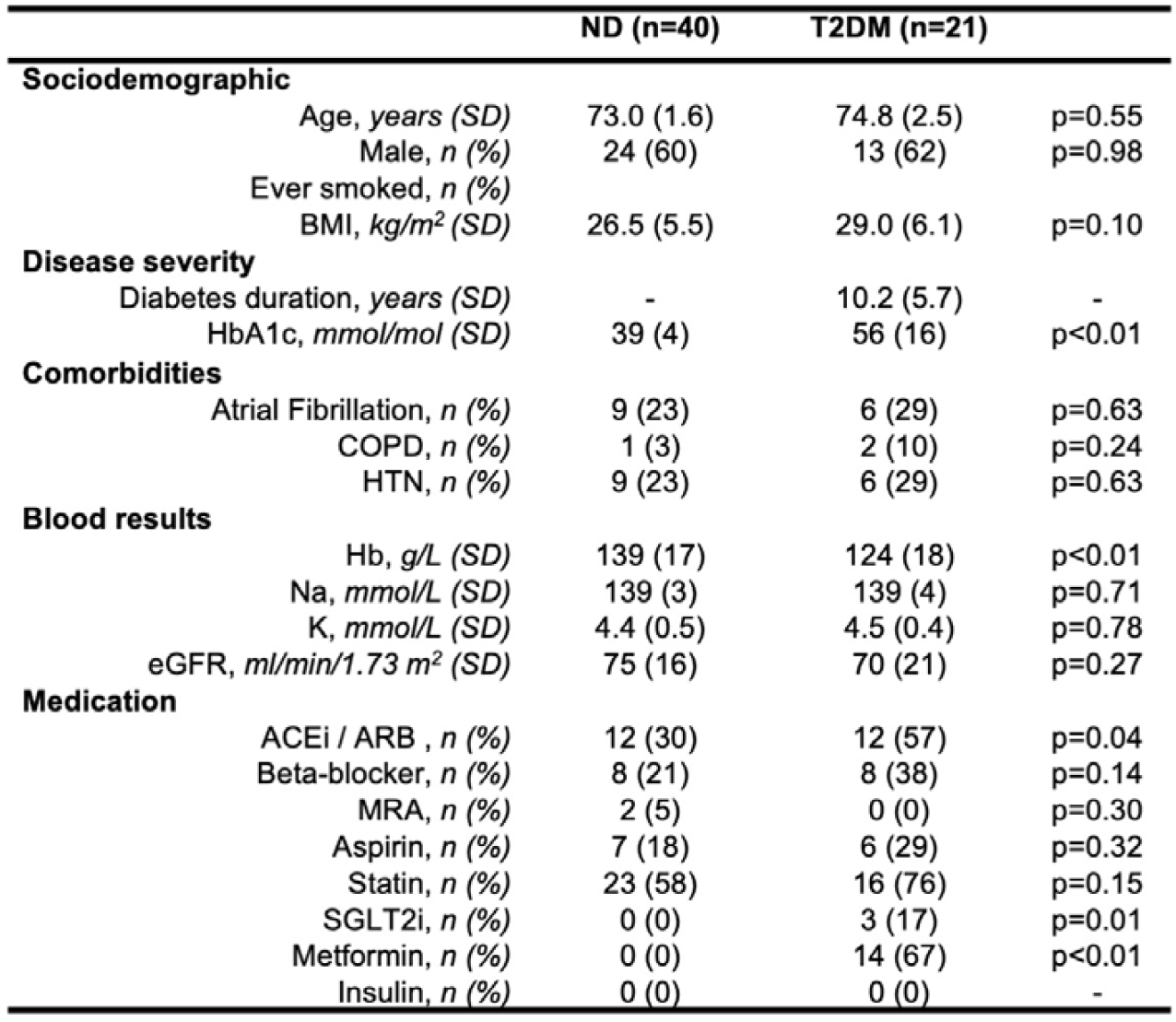
Characteristics of study participants. **a**, Distribution of sociodemographic, disease severity, comorbidities, blood results, and medication criteria of ND (n=40) and T2DM (n=21) individuals participating in this study.

## Supplementary material and methods

### Ethics declarations

The United Kingdom Health Research Authority provided ethical approval for the study following review by the Leeds West Research Ethics Committee (11/YH/0291). All patients provided written informed consent and the study was conducted in accordance with the principals of the Declaration of Helsinki.

### Recruitment methodology

Patients with and without T2DM undergoing cardiac device implantation at the Leeds Teaching Hospitals Trust were recruited between 2018 to 2025. For inclusion patients were required to be prescribed anti-diabetic medicine and/or have a serum HbA1c ≥48mmol/mol. We also required all patients to have normal left ventricular systolic function (left ventricular ejection fraction ≥55%).

### Tissue sampling

At the time of cardiac device implantation and under local anaesthetic, a small (approximately 1-1.5cm^3^ or 250-500mg) sample of subcutaneous adipose tissue was taken from the area superficial to pectoralis major and a small piece (approximately 1-1.5cm^3^ or 250-500mg) of skeletal muscle was harvested from the pectoralis major. The tissue samples were immediately placed into MACS tissue storage solution for endothelial cell extraction. Blood samples were taken at the time of procedure through the pacing lead sheath for routine biochemistry and haematology (including serum NT-proBNP and HbA1c).

### Bulk RNA transcriptomics sequencing and gene expression analysis

RNA sequencing (RNA-seq) count data were collected from multiple independent experimental batches. To ensure consistent annotation, count matrices were merged into a unified dataset. Because the dataset was originated from multiple sequencing runs, batch effect was explicitly modelled as a covariate in the DESeq2 design formula (∼batch + condition). This approach ensured that the technical variation attributable to sequencing batches was minimised while biological differences between conditions were preserved. For the presented datasets, we only adjusted for batch effect.

Differential expression analysis was performed using the DESeq2 package. Genes with fewer than 10 read counts across total samples were excluded to reduce noise in the analysis. Negative binomial generalised linear models were fitted to each gene, with dispersion estimates calculated. Wald tests were used to identify differential expression between conditions. P-values were adjusted using the Benjamini–Hochberg procedure to control the false discovery rate (FDR). Genes with adjusted p-values < 0.05 were considered significantly differentially expressed.

### GO-term analysis

GO-term analysis was performed with g:Profiler http://biit.cs.ut.ee/gprofiler/ using Benjamini-Hochberg FDR and a user-defined p-value threshold of 0.05.

### Cell isolation and culture

Both MVEC from pectoral SAT and SkM tissue were isolated and maintained as described previously (5). Briefly, tissue was digested using 0.1U/ mL collagenase/ 0.8U/ mL dispase (Roche), dead cells were removed using the dead cell removal kit on a LS column, and live cells were purified using anti-CD31 coated magnetic beads on a MS column; all materials Milltenyi Biotec. Cells were routinely assessed for purity using flow cytometry, achieving ∼99% purity on average. Cells were maintained in ECGM-MV medium (Promocell). Cells were routinely frozen in Cryo-SFM (Promocell). After defrosting, all cells were passaged once. Unless otherwise stated, cells were seeded at 5×10^3^/cm^2^ and grown to confluency. Cells not reaching confluency within seven days after seeding were discarded.

### Morphometrics

SAT and SkM MVECs were subjected to label free morphological profiling through brightfield imaging using a 4L BRVY Attune Cytpix Flow Cytometer (Thermo Fisher Scientific) integrating acoustic focusing with high-speed brightfield imaging for acquisition of morphological data using appropriate FSC and SSC voltages to capture all samples. Brightfield images were processed with cell full resolution settings according to manufacturer’s recommendation prior to dimension reduction. Dimension reduction with automatic clustering was performed to identify maximum clustering and assessment of quantitative morphological features. Unsupervised clustering was applied to identify phenotypically distinct subpopulations. The dimensionality reduction technique ‘uniform manifold approximation and projection’ (UMAP) was employed for visualization. Data analysis was be performed using GraphPad Prism, Attune Cytometric Software v7.1 and Anaconda. Comparisons between groups were performed using appropriate statistical tests.

### Western blot analysis

Western blotting was performed as described in (19). Briefly, cells were grown to confluency, washed twice in M199 (Gibco) supplemented 10 mM HEPES and 1 mM Sodium Pyruvate, and incubated for four hours in ECGM-MV (Promocell). Cells were washed once in ice-cold PBS prior to lysis in cell extraction buffer (Invitrogen). Protein concentration was determined using a BCA protein assay kit (Pierce). 15 µg protein was loaded onto 4—12% NuPAGE Bis-Tris mini protein gels (Invitrogen) and subsequently transferred onto a nitrocellulose membrane using the Trans-Blot Turbo system (BioRad). Non-specific binding of antibodies was blocked by using 3% BSA/ TBS-T (Tis-buffered saline, 0.1% Tween-20) for 30-60 min. Primary antibody incubation was performed over-night at 4°C followed by four 5 min washes in TBS-T using the following dilutions: VCAM-1 (Abcam, ab174279, 0.1 mg/mL); eNOS (BD Biosciences, 610297, 0.2 mg/mL); cleaved caspase-3 (9661, 1:1000), peIF2a (3398, 1:1000), eIF2a (5324, 1:1000), IL-6 (12153, 1:1000), IL-8 (94407, 1:1000), MCP-1 (39091, 1:1000), PAI-1 (49536, 1:1000), p16 (80772, 1:1000), p21 (2947, 1:1000), TNF-a (6945, 1:1000), all Cell Signaling Technology; LC3B (GTX127375, 0.5 mg/mL), p62 (GTX629890, 0.5 mg/mL), all GeneTex; ATF6 (NBP1-40256, 1 mg/mL), BIP/ GRP78 (NBP1-06277, 0.5 mg/mL), CHOP/ GADD153 (NBP1-06277, 1 mg/mL), pIRE1a (NB100-2323, 1 mg/mL), IRE1a (NB100-2324, 1 mg/mL), all Novus Biologicals; a-SMA (R&D Sytems, MAB1420, 0.2 mg/mL); HSP90 (sc-13119, 4 ng/mL; used as loading control), ICAM-1 (sc-107, 0.2 mg/mL), VE-cadherin (sc-9989, 0.2 mg/mL), all Santa Cruz Biotechnology. Horseradish peroxidase (HRP)-conjugated antibodies (Cytiva) were used for 60 min followed by four 5 min washes in TBS-T. Image acquisition was performed on a Syngene G:Box GTX4 system using enhanced chemiluminescence (Millipore). Analysis was performed using FIJI ImageJ.

### ELISA

Serum C-peptide levels were determined with a human C-peptide ELISA (R&D Systems, DICP00) according to the manufacturer’s directions.

### Flow cytometry

Purity assessment of cultured MVEC was performed using a Beckman Coulter CytoflexS Flow Cytometer using the CytExpert software (Beckman) for analysis. Cells were labelled with the following Milltenyi Biotec antibodies at a 1:50 dilution: CD45-FITC (130-110-631), CD31-PerCP-Vio700 (130-110-673), CD144 (VE-Cadherin)-PE (130-118-358).

### Total RNA extraction and RT-qPCR

Total RNA was extracted from confluent cells using TRIzol reagent (Invitrogen) following the manufacturer’s instructions. RNA Clean & Concentrator-5, (Zymo Research) was used for further RNA purification. Extracted RNA was analysed using a NanoDrop spectrophotometer (Thermo Fisher Scientific) for quantity and quality. Reverse transcription was performed using Lunascript RT SuperMix (NEB) following the manufacturer’s instructions on an AB Veriti 96-well Thermal Cycler (Applied Biosystems). RT-qPCR was performed using iTaq Universal SYBR Green Supermix (BioRad) using 15 ng of cDNA per well was on a LightCycler 96 (Roche) following the manufacturer’s suggestions. Gene expression was normalised to GAPDH, mRNA expression values were calculated using the 2^-ΔΔCt^ method. For each cell type, data from T2DM patients was compared to that of the non-diabetic control patients and expressed as fold difference. All primers were purchased from BioRad: CD44 (qHsaCID0013679), GAPDH (qHsaCED0038674), SNAI1 (qHsaCED0057267), vWF (qHsaCED0033955), and ZEB2 (qHsaCED0038149).

### Senescence-associated β-galactosidase assay

For detection of senescence associated β-galactosidase, the CellEvent Senescence Green Detection kit (Invitrogen) was used according to the manufacturer’s suggestions. SAT MVEC and SkM MVEC were seeded to confluency into six replicate wells of a 96-well plate (Greiner) at 3×10^4^ cells/ cm^2^ 24 h prior to experimentation. Cells were imaged on the EVOS imaging system (Thermo-Fisher). For fluorescence analysis, the triangle threshold algorithm at auto-setting in FIJI ImageJ was employed.

### Statistical methodology

Statistical analysis was performed using GraphPad Prism version 10 or R. All data were tested for normality and equal variance followed by the suitable statistical test. Where appropriate, individual data points are shown. Sample sizes were not pre-determined. Linear regression models were constructed for CHOP and clinical characteristics adjusting for age and biological sex. All statistical tests were two-sided, and statistical significance was defined as p > 0.05. Missing data were not imputed.

## Notes

### Competing Interest Statement

The authors have declared no competing interest.

